# An integrated semantic publishing workflow for taxonomy: a review of the Afrotropical dung beetle genus *Amietina* (Coleoptera: Scarabaeinae)

**DOI:** 10.64898/2026.10.19.760055

**Authors:** Giulio Montanaro, Neema Kilimba, Wilfred Sengo, Sergei Tarasov

**Affiliations:** Finnish Museum of Natural History, Pohjoinen Rautatiekatu 13, FI-00014 Helsinki, Finland; Tanzania Wildlife Research Institute (TAWIRI), P.O. Box 661, 206 Njiro Road, 2113 Lemara, Arusha, Tanzania

**Author notes:** Corresponding author: Giulio Montanaro.

**Keywords:** LaTex, collaborative writing, FAIR principles, new species, Onthophagini, Eastern Arc Mountains, Tanzania

## Abstract

Taxonomic studies generate large amount of descriptive information, yet species treatments are usually written directly as natural-language text. This approach makes the underlying phenotypic information difficult to extract, reuse computationally, and maintain consistently across studies. We present an integrated workflow connecting Phenoscript, GitHub, and Overleaf. Species descriptions are written semantically in Phenoscript using biological ontologies, making their phenotypic content computable and reusable; they are then stored and version-controlled in GitHub and synchronized with an Overleaf manuscript, where human-readable descriptions are rendered directly for publication. The workflow reduces repetitive copying and formatting, preserves revision history, and promotes the production of FAIR phenotypic and taxonomic data.

We apply this workflow to the African dung beetle genus *Amietina* Cambefort, 1981 (Coleoptera: Scarabaeinae: Onthophagini), and describe four new species from Tanzania—*A. bondarevi* **n. sp**., *A. grebennikovi* **n. sp**., *A. mazumbaiensis* **n. sp**., and *A. praedicta* **n. sp**.—bringing the total number of recognized species to eight. We also provide new distributional records, present an updated identification key, and discuss microendemism in *Amietina* across the Eastern Arc Mountains.

## INTRODUCTION

Taxonomic revisions turn observations into species descriptions, diagnoses, identification keys, and nomenclatural acts. Yet their preparation is still largely document-based: observations are rewritten as prose, edited in a manuscript, and reformatted for publication, typically in Microsoft Word or Google Docs. Repeatedly moving information between data and text takes time, makes it harder to keep them consistent, and can introduce errors. It also leaves much of the information in taxonomic publications difficult to find and reuse (Smith et al., 2013; Miller et al., 2012; Penev et al., 2017).

This problem is especially clear in species descriptions. Natural language is easy for taxonomists to read, but computers cannot reliably identify the anatomical characters and their relationships described in the text. The same structure may also be named differently by different authors, while the meaning of a term may depend on the taxonomic group or context. These observations must therefore be extracted and recoded before they can be searched, compared across taxa, or combined with other data.

Semantic descriptions based on ontologies address this limitation by representing observations with standardized terms and explicit relationships (Balhoff et al., 2013; Deans et al., 2015). In simple terms, an ontology provides agreed names and definitions for biological terms and specifies how they are related.

Each term has a stable identifier, allowing descriptions that use the same concept to be linked even when their wording differs. Phenoscript provides a formal language for writing such descriptions and can generate both machine-readable data and human-readable text for a conventional taxonomic treatment (Montanaro and Tarasov, 2024; Montanaro et al., 2024a). Taxonomists can therefore read the resulting description in a familiar form, while its underlying content remains available for computational searches and comparisons.

Generating readable text, however, does not solve the problem if it must be copied into the manuscript again after every change. We therefore connect Phenoscript, GitHub, and Overleaf in a single workflow to streamline this process. Phenoscript is the source of the species descriptions, Git records changes to the descriptions and related files, and Overleaf compiles the LATEX manuscript containing the generated text. The descriptions are written once in Phenoscript and rendered directly in the manuscript. When a description is corrected, its output can be regenerated and synchronized through GitHub instead of being copied, reformatted, and checked again by hand. This streamlines the manuscript preparation and reduces the risk that the published treatment differs from its semantic source.

The same connection also makes the wider manuscript easier to manage. GitHub keeps the descriptions, manuscript, bibliography, figures, and processing files together and preserves a history of their changes. Collaborators can see what was changed, compare versions, and recover earlier content. Overleaf provides a shared writing environment and automatically compiles these files into the formatted paper. This approach keeps the manuscript closer to its source data and makes collaboration more transparent and reproducible (Ram, 2013; Braga et al., 2023; Himmelstein et al., 2019). Because LATEX separates content from formatting, the same source can also be adapted to journal templates with less manual reformatting (Bar and Wang, 2021; Krewinkel and Winkler, 2017).

Here, we use this workflow in a taxonomic review of the Afrotropical dung beetle genus *Amietina* Cambefort, 1981 (Coleoptera: Scarabaeinae: Onthophagini). The genus currently contains four described species from forests in West, Central, and southeastern Africa (Cambefort, 1981; Branco, 1989). Material collected in the Eastern Arc Mountains of Tanzania revealed four additional species, broadening the morphological limits of the genus, which is here redefined. The discovery of these species also add to the limited knowledge of dung beetle endemism across the Eastern Arc mountain blocks (Burgess et al., 2007; Montanaro et al., 2024b). Herein, we describe *A. bondarevi* **n. sp**., *A. grebennikovi* **n. sp**., *A. mazumbaiensis* **n. sp**., and *A. praedicta* **n. sp**. as well as the previously unknown female of *A. nyassalandica*. Additonally, we provide new distributional records, and present an updated identification key. Alongside these taxonomic contributions, we show how Phenoscript, GitHub, and Overleaf can connect semantic phenotype data directly to a publication-ready manuscript.

This manuscript conforms to the requirements of the amended International Code of Zoological Nomenclature (ICZN). Please note that this work is not issued for the purpose of zoological nomenclature and it should be considered not published within the meaning of the code (ICZN code, Section 8.2). The nomenclatural acts and new taxon names described herein should therefore be considered as not available.

## MATERIALS AND METHODS

### Workflow integrating Phenoscript, GitHub, and Overleaf

We organized the project as a four-step workflow linking semantic description, version control, collaborative writing, and manuscript rendering (Fig. 1). The GitHub repository contains the Phenoscript source files, generated outputs, manuscript source, bibliography, and figures, thereby keeping the material required to reproduce the taxonomic treatments in one version-controlled project.

**Figure 1.**
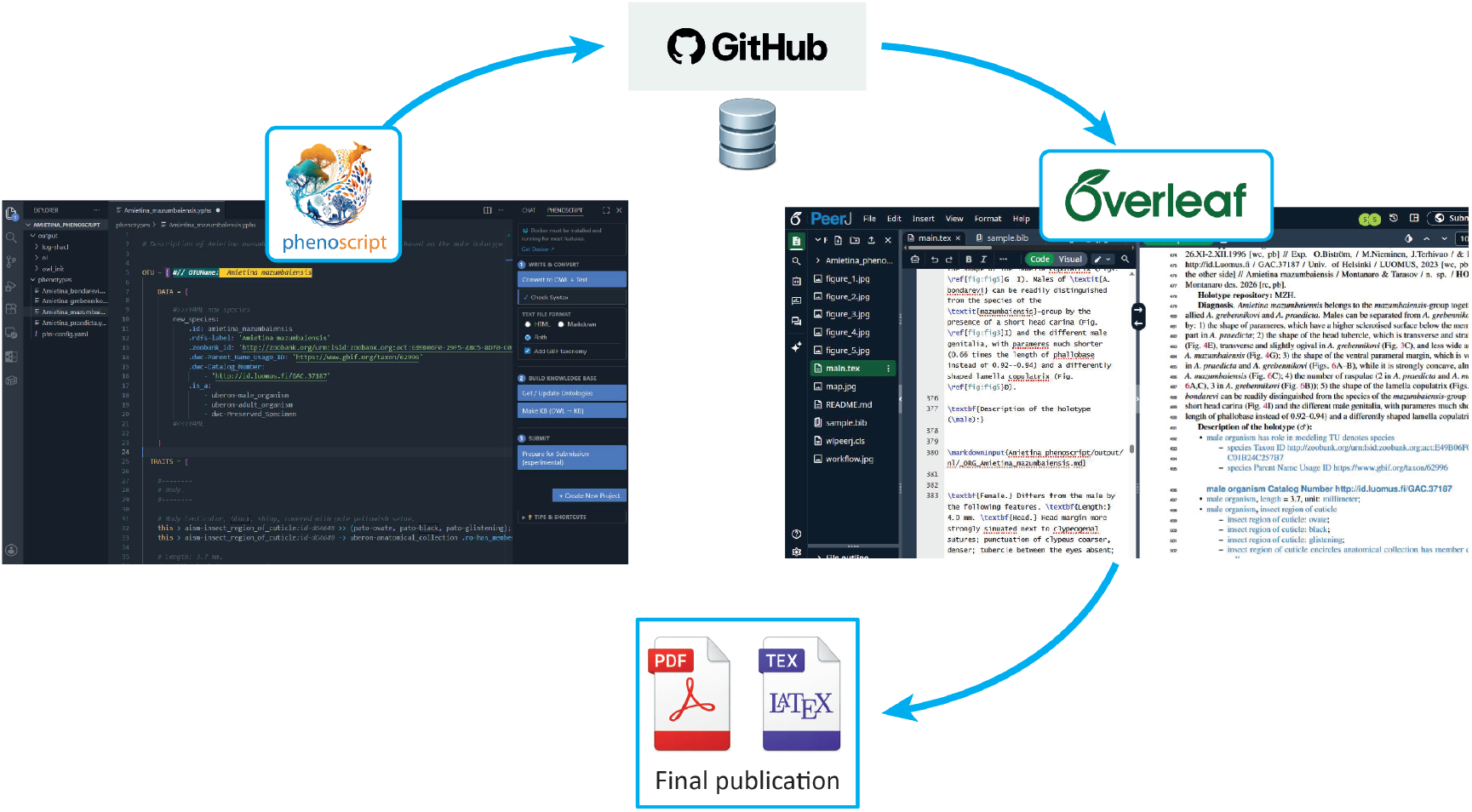
Workflow used to prepare this publication. Taxonomic descriptions were written in Phenoscript, validated, and converted into human-readable Markdown and machine-readable OWL formats. The Phenoscript source files and generated outputs were committed to a GitHub repository together with the manuscript, bibliography, and figures. The manuscript was prepared in LATEX using Overleaf and synchronized with the GitHub repository, allowing the generated Markdown descriptions to be rendered directly within the taxonomic treatments

**Figure 2.**
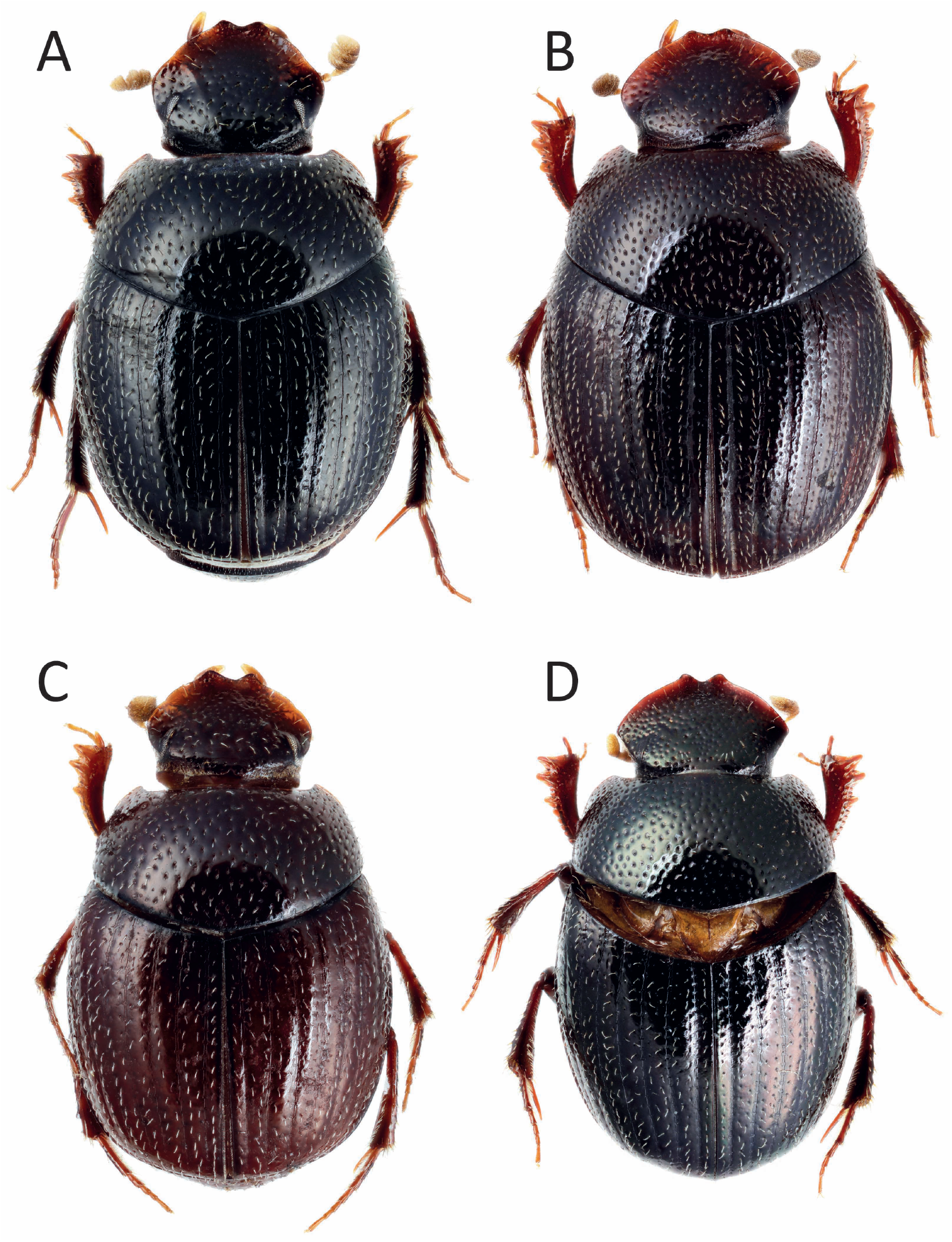
*Amietina* species of the *larrochei*-group, habiti: *A. larrochei*, male (A); *A. eburnea*, female (B); *A. cambeforti*, male (C); *A. nyassalandica*, female (D).

**Figure 3.**
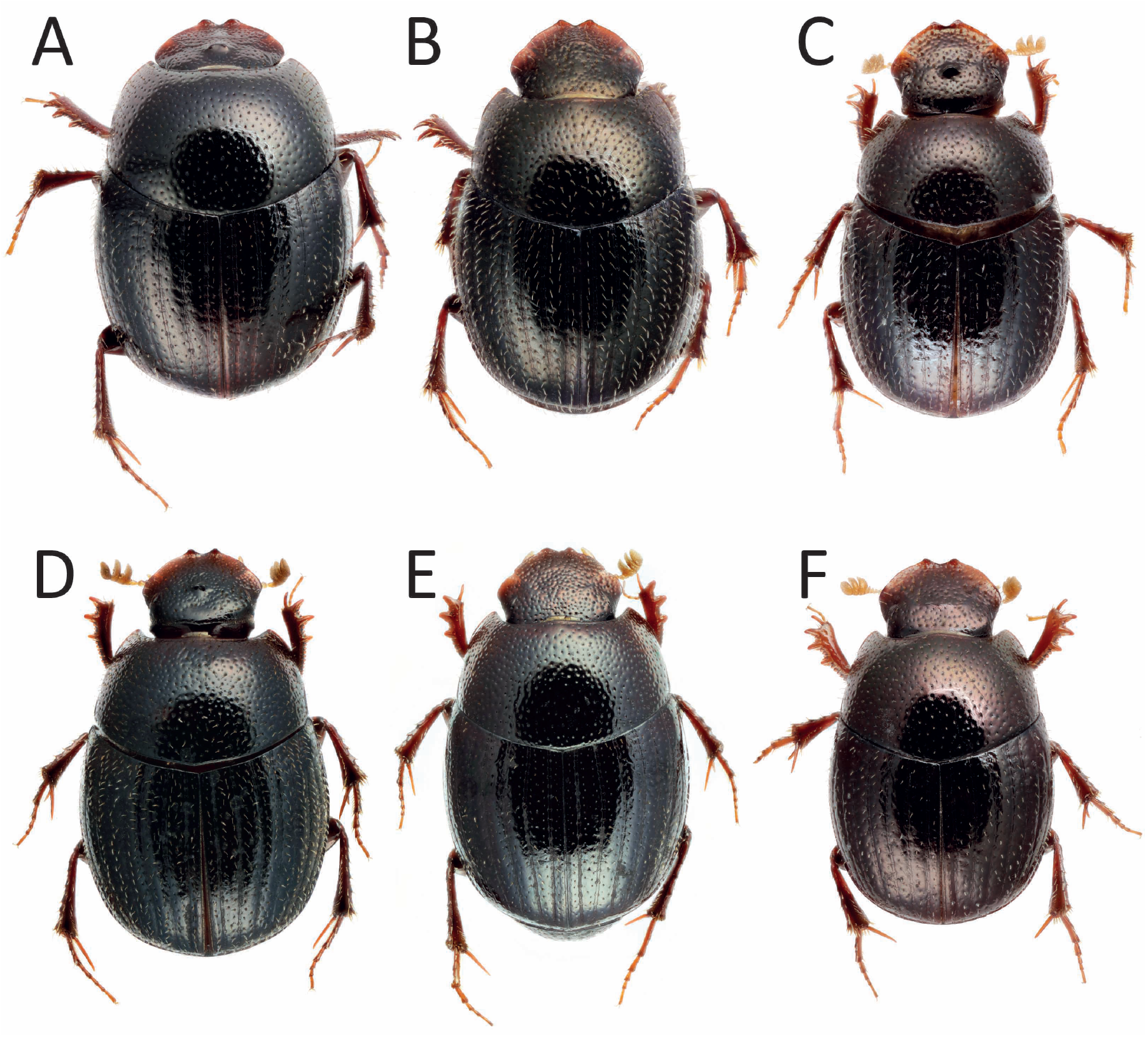
*Amietina* species from Tanzania, habiti: *A. praedicta* **n. sp**., male (holotype) (A) and female (paratype) (B); *A. grebennikovi* **n. sp**., male (holotype) (C); *A. mazumbaiensis* **n. sp**., male (holotype) (D) and female (paratype) (E); *A. bondarevi* **n. sp**., male (holotype) (F).

**Step 1: Writing and processing semantic descriptions**. We wrote the descriptions of the four new species as .yphs files in Visual Studio Code, using the dedicated Phenoscript extension. The Phenoscript workflow parsed each description into an ontology-based knowledge graph, checked it against the project validation rules, and generated machine-readable OWL and human-readable Markdown and HTML outputs (Montanaro and Tarasov, 2024; Montanaro et al., 2024a). Thus, the semantic source and the text used in the taxonomic treatment were produced from the same description.

**Step 2: Preparing the manuscript**. We wrote the manuscript in LATEX using Overleaf. Overleaf provided a shared editing environment and compiled the manuscript, bibliography, figures, cross-references, and imported descriptions into a formatted PDF. Because formatting was controlled by the LATEX document class and commands, changes to the journal style did not require manual reformatting of each taxonomic treatment.

**Step 3: Versioning and synchronizing the project**. We used GitHub as the link between the local Phenoscript project and the Overleaf project. Changes made locally were committed and pushed to the GitHub repository, which preserved the revision history of both the semantic descriptions and their generated outputs. The repository was then synchronized with Overleaf through its GitHub integration. This arrangement allowed collaborators to compare revisions, recover earlier versions, and update the manuscript and its source data within the same project (Ram, 2013; Braga et al., 2023).

**Step 4: Rendering descriptions in the manuscript**. We inserted each generated Markdown description directly into the corresponding taxonomic treatment with the \markdownInput{} command from the markdown package. After a Phenoscript source file was revised, we regenerated its outputs, committed them to GitHub, and synchronized the repository with Overleaf. The next manuscript compilation then rendered the revised description without manual copying or Markdown-to-LATEX conversion. The Phenoscript file therefore remained the authoritative source for both the computable phenotype data and the description read in the manuscript.

### Semantic descriptions using Phenoscript

We prepared separate Phenoscript descriptions for *Amietina bondarevi, A. grebennikovi, A. mazumbaiensis*, and *A. praedicta*, following the approach of Montanaro and Tarasov (2024) and Montanaro et al. (2024a). To keep descriptions consistent among species, we used the same order of anatomical regions and followed the project annotation guide. Comments in the .yphs files record the corresponding conventional wording and make the semantic statements easier to review.

Each statement links an anatomical entity to a quality via relationship. Anatomical terms were drawn primarily from the Anatomy Ontology of Insect Skeleto-Muscular system (AISM) and the Coleoptera Anatomy Ontology (COLAO; Girón et al., 2024). Phenotypic qualities were represented with terms from the Phenotype and Trait Ontology (PATO), spatial regions with the Biological Spatial Ontology (BSPO), relations with the Relations Ontology (RO), and measurement units with the Units of Measurement Ontology (UO).

We processed each source description to produce an OWL representation of the phenotype and checked the output against the validation rules implemented in the Phenoscript workflow. The final Markdown files present the same statements in a readable form and retain hyperlinks from anatomical and phenotypic terms to their ontology records. These Markdown files, rather than separately written descriptions, are the files rendered in the four new-species treatments below.

### Material examined and morphology

Male and female specimens of all described *Amietina* species were examined, except for *A. nyassalandica* of which we studied a single female, while male characters were retrieved from the detailed original description by Branco (1989). Although we did not examine type material of any of the species, we studied specimens of *A. cambeforti, A. eburnea* and *A. larrochei* coming from the same collecting events of those examined by Branco (1989) and on whose correct identification we have no doubts. Field work and collection of specimens in Tanzania was authorised by the Tanzania Wildlife Research Institute (TAWIRI) (clearance AB.235/325/01”A”/61) and the Tanzania Commission for Science and Technology (COSTECH) (permit CST00001334-2025-2025-00982). Studied specimens are preserved in the following collections:

MZHF–Finnish Museum of Natural History LUOMUS, Helsinki, Finland;

GMOC–Giulio Montanaro personal collection, Creazzo, Italy;

JFJC–Jean-François Josso personal collection, Muzillac, France;

An additional specimen from the Rubeho Mountains putatively attributable to *Amietina* was identified among the samples registered in the Barcode of Life Database (BOLD) (unique ID: CNCCOLVG00004064) by the colleague Vasily Grebennikov (Canadian Food Inspection Agency, Ottawa, Canada). However, despite the kind efforts of V. Grebennikov and Bruce Gill (Canadian National Collection of Insects, Ottawa, Canada), it was not possible to locate the physical voucher, which shall be considered lost. We therefore do not treat that specimen here.

According to the identification key by d’Orbigny (1913) on African Onthophagini, the new species could belong to the *Onthophagus* species-groups 3 (females) or 19 (males; dichotomy 41(34)). The comparison with the original descriptions of all valid species of those groups did not result in any correspondence.

Morphological terminology and protocols follow Montanaro et al. (2024a) and Tarasov and Solodovnikov (2011). For male genitalia, the length of the phallobase was measured from the middle of the antero-dorsal bulge to the middle of the postero-dorsal margin; parameral length was measured from the postero-dorsal angle to the apex (*i*.*e*., the distalmost part) of the left paramere. Pictures of morphology were taken with a Canon EOS 5DS and those of genitalia with a Canon EOS 250D connected to a stereomicroscope. The distribution map was realised in QGIS (QGIS Development Team, 2009) and Eastern Arc forests outlined using the Terrestrial Ecoregions of the World layer (Olson et al., 2001).

For holotype labelling, we use “//” to separate different labels and “/” to separate lines on the same label. Comments to labels are given between square brackets. The following abbreviations are used: wc (white cardboard), rc (red cardboard), hb (handwritten in black), pb (printed in black).

## TAXONOMY

### *Amietina* Cambefort, 1981 *sensu novo*

Cambefort (1981): 143 (original description);

Cambefort (1984): 106 (ecology);

Cambefort (1985): 339 (faunistics);

Branco (1989): 201 (description of 2 new species);

Nummelin and Hanski (1989): 351 (ecology);

Cambefort and Walter (1991): 200 (ecology);

Cambefort and Bordat (2003): 559 (faunistics);

Davis et al. (2008): 216 (systematics, natural history);

Josso and Prévost (2008): 2 (systematics);

Moretto (2010a): 469 (faunistics);

Moretto (2010b): 14 (ecology, natural history);

Branco (2011): 11 (nomenclature);

Philips et al. (2023): 2 (ecology).

**Type species**: *Amietina larrochei* Cambefort, 1981 (by original designation).

#### Diagnosis

The genus is a member of the tribe Onthophagini. It can be differentiated from all other genera and subgenera of *Onthophagus* by the following combination of characters: 1) body small (2.7–4.1 mm), lenticular; 2) antennae with 8 segments; 3) clypeus with two teeth; 4) protibiae with 3 teeth on dorsal margin (Figs. 4J—O); 5) male protibiae with a small, sharp uncus (Figs. 4J,L,M,O); 6) dorsal body surface covered with simple punctures bearing conspicuous setae; 7) parameral apices acuminate and bent ventrally (Figs. 5–6); 8) lamella copulatrix composed by two adjacent but separate endophallites, usually with an additional, elongated small distal endophallite (Figs. 6E–I). The following two characters, previously used to define the genus, are here reconsidered: 1) head surface either devoid of protrusions (but sometimes with a transverse bulging on the frons) (Figs. 4A–D) or with a small frontal tubercle or short carina (Figs. 4E–I); 2) distal protibial margin from perpendicularly to obliquely truncated (Figs. 4J–O).

**Figure 4.**
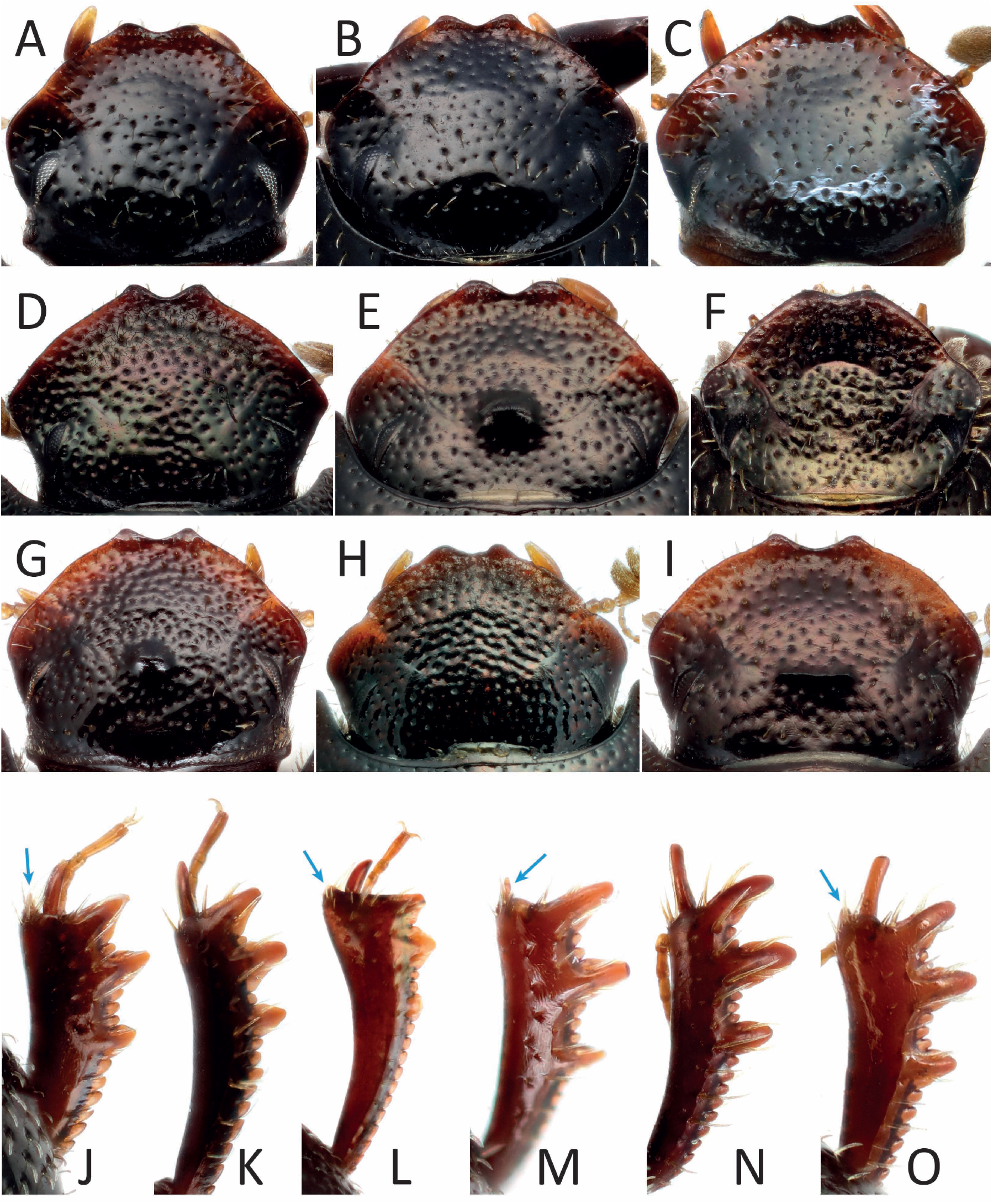
Heads and protibiae of *Amietina* species. Heads in dorsal view: *A. larrochei*, male (A) and female (B); *A. eburnea*, male (C); *A. nyassalandica*, female (D); *A. praedicta* **n. sp**., male (holotype) (E) and female (paratype) (F); *A. mazumbaiensis* **n. sp**., male (holotype) (G) and female (paratype) (H); *A. bondarevi* **n. sp**., male (holotype) (I). Protibiae in dorsal view, blue arrows pointing at unci: *A. larrochei*, male (J) and female (K); *A. eburnea*, male (L); *A. mazumbaiensis* **n. sp**., male (holotype) (M); *A. praedicta* **n. sp**., female (paratype) (N); *A. bondarevi* **n. sp**., male (holotype) (O).

**Figure 5.**
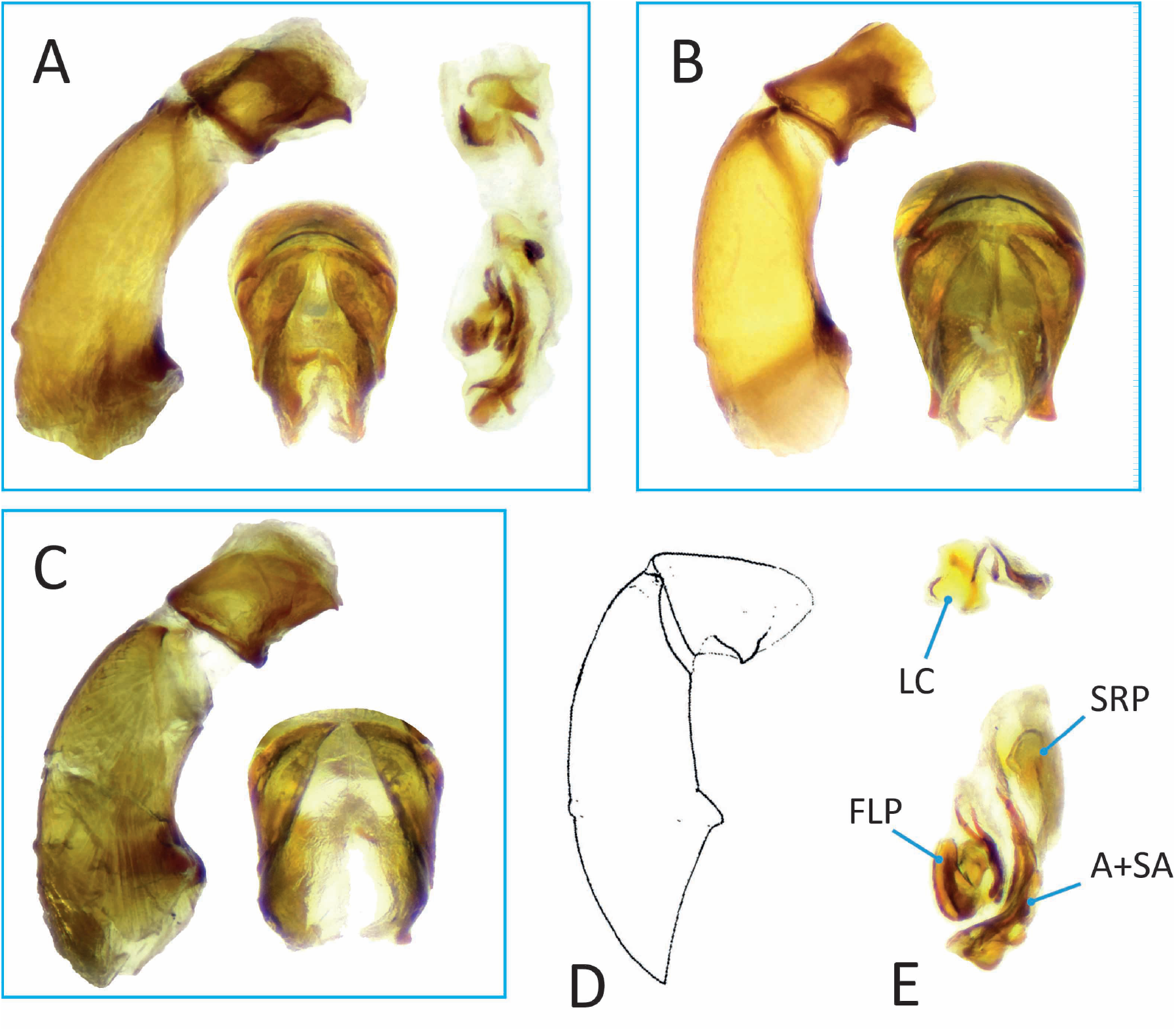
Male genitalia of *Amietina* species of the *larrochei*-group. *A. larrochei* (A): aedeagus in lateral (left) and dorsal (middle) views, and endophallus (right); *A. eburnea* (B): aedeagus in lateral (left) and dorsal (right) views; A. cambeforti (C): aedeagus in lateral (left) and dorsal (right) views; *A. nyassalandica* (D), aedeagus from Branco (1989); *A. eburnea* (E), endophallus illustrating endophallites: lamella copulatrix (LC), frontolateral peripheral (FLP), superior right peripheral (SRP), axial and subaxial (A+SA).

**Figure 6.**
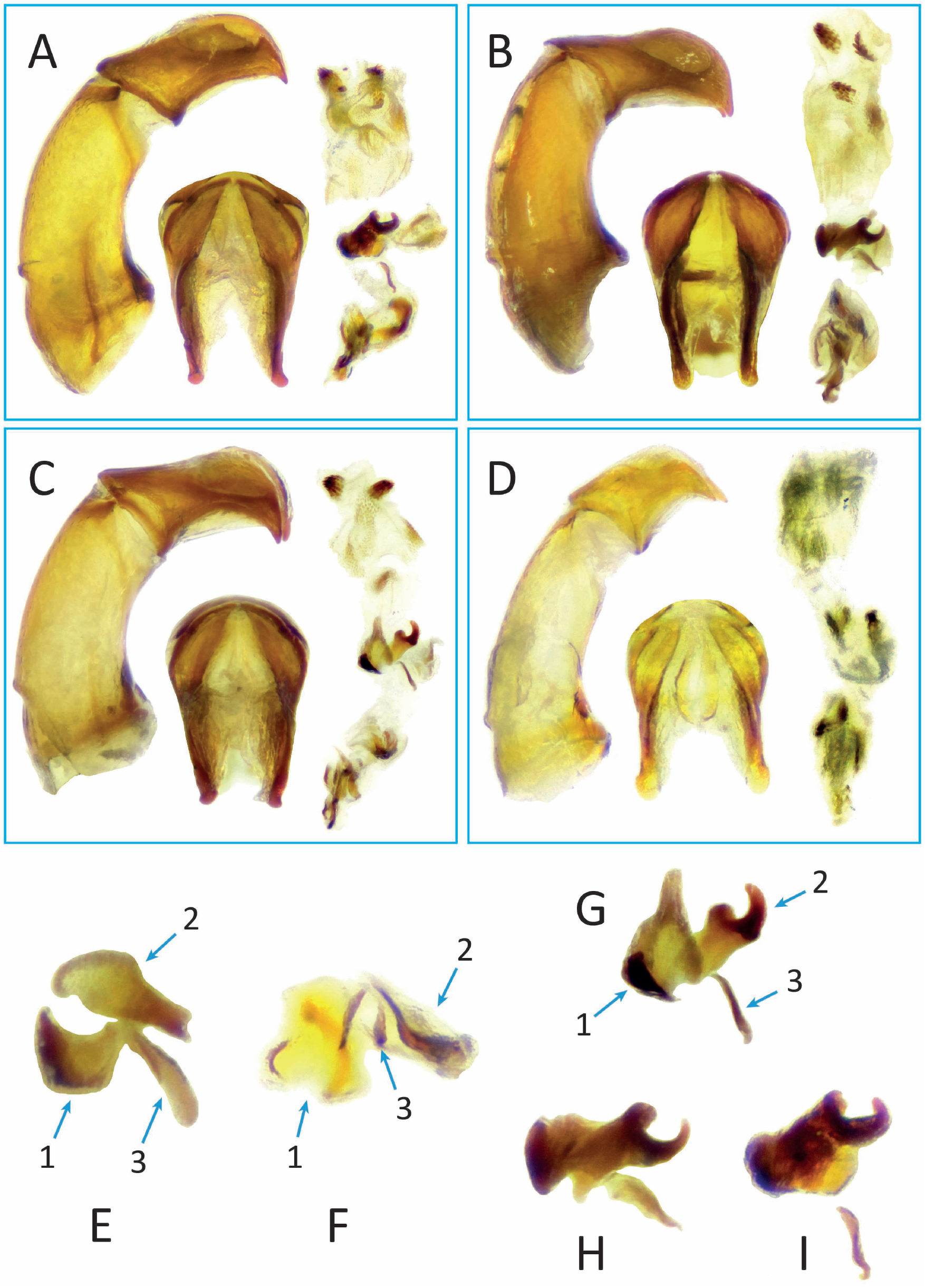
Male genitalia of *Amietina* species. Genitalia of holotypes of the *mazumbaiensis*- and *bondarevi*-groups, showing aedeagus in lateral (left) and dorsal (middle) views, and endophallus (right) (A–D): *A. praedicta* n. sp. (A); *A. grebennikovi* **n. sp**. (B); *A. mazumbaiensis* **n. sp**. (C); *A. bondarevi* **n. sp**. (D). Lamellae copulatrices of Amietina species, showing the three endophallites (1–3) composing them: *A. larrochei* (E), *A. eburnea* (F), *A. mazumbaiensis* (G), *A. grebennikovi* (H) and *A. praedicta* (I).

**Figure 7.**
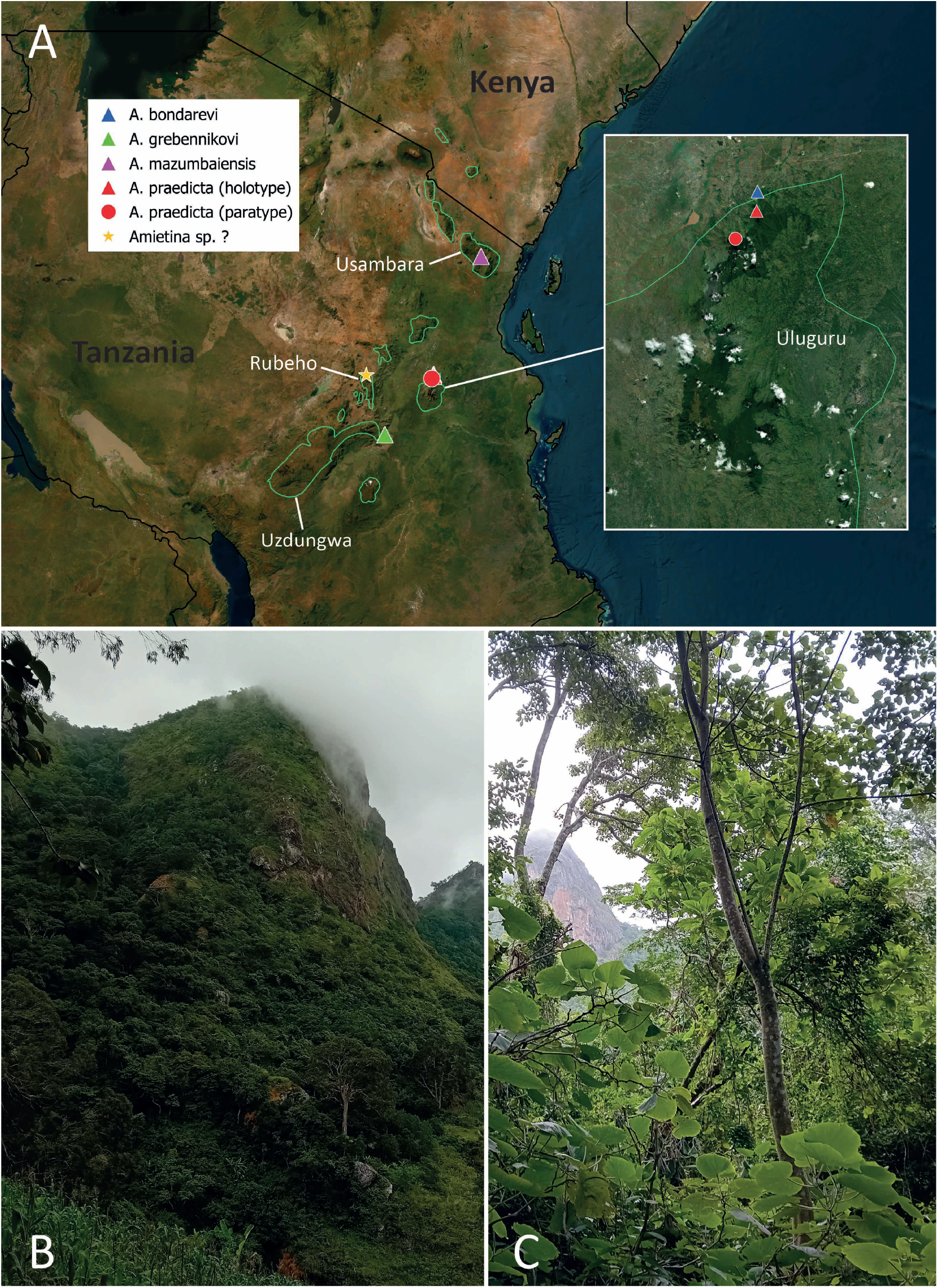
Distribution and habitat of *Amietina* species. Map showing the distribution of *A. bondarevi* n. sp., *A. grebennikovi* n. sp., *A. mazumbaiensis n. sp*. and *A. praedicta* n. sp. in Tanzania, with the boundaries of the Eastern Arc Mountains ecoregion highlighted in green (A); rainforest edge (B) and vegetation (C) at the foothills of the northern Uluguru Mountains close to the Lupanga Peak, type locality of *A. praedicta* (pictures by G. Montanaro).

#### Biology

Based on literature (Branco, 1989) and the present data, *Amietina* species are forest-specialised. Specimens were collected in ant nest debris, excrements, rotten fish, fruits and dead diplopods, suggesting a rather generalist feeding behaviour (present data; Branco, 1989; Cambefort, 1981; Moretto, 2010b).

#### Distribution

The genus is endemic to the Afrotropical region.

#### Remarks

*Amietina sensu novo* maintains a substantially similar definition to its previous concept (Branco, 1989; Cambefort, 1981). The only differences are that the distal margin of protibiae can be either perpendicularly (as in the previous concept, except for *A. larrochei*) or obliquely truncated (new concept), and that the head can be either unarmed (previous concept) or with protrusions (new concept). Furthermore, *Amietina* species are characterised by a lamella copulatrix formed by 3 separate sclerites characteristically arranged (Figs. 6E–I). The only exception is *A. bondarevi*, which has only 2 evident sclerites, but this may be due to the poor sclerification of the endophallus of the only examined specimen, which may have rendered the third, smaller sclerite less clearly visible.

Based on morphology, the genus can be subdivided into three operational species-groups:

1. *Amietina larrochei* species-group, characterized by: 1) head without protrusions in both sexes, surface at most swollen medially (Figs. 4A–D); 2) distal margin of male protibiae perpendicularly truncated or almost so (Figs. 4J,L); 3) proximal region of endophallus without raspulae (Figs. 5A,E); 4) pronotal punctures with uniform to irregular size, occasionally with some distinctly smaller punctures intermixed with bigger ones (*A. larrochei*) or with micropunctuation (*A. nyassalandica*) (Fig. 2). The group comprises *A. cambeforti* Branco, 1989, *A. eburnea* Cambefort, 1981, *A. larrochei* Cambefort, 1981 and, putatively, as we could not examine male genitalia, *A. nyassalandica* Branco, 1989.
2. *Amietina mazumbaiensis* species-group, characterized by: 1) head with a tubercle between the anterior margin of the eyes (males) or a short, arched carina on the frons (females) (Figs. 4E–H); 2) distal margin of male and female protibiae obliquely truncated (Figs. 4M–N); 3) proximal region of endophallus with raspulae (Figs. 6A–C); 4) pronotal punctuation clearly heterogeneous, composed of larger punctures intermixed with many smaller ones. The group comprises *A. grebennikovi* **n. sp**., *A. mazumbaiensis* **n. sp**. and *A. praedicta* **n. sp**.
3. *Amietina bondarevi* species-group, characterized by: 1) head with a short carina between the anterior margin of the eyes (Fig. 4I); 2) distal edge of protibiae obliquely truncated (Fig. 4O); 3) proximal region of endophallus without raspulae (Fig. 6D); 4) pronotal punctuation clearly heterogeneous, composed of larger punctures intermixed with many smaller ones. The group comprises the only *A. bondarevi* **n. sp**.

Other undescribed species may also belong to the genus as defined here (Moretto, 2010b; P. Moretto, personal communication) (see also the Discussion).

### *Amietina cambeforti* Branco, 1989

(Figs. 2C, 5C)

Branco (1989): 201 (original description);

Nummelin and Hanski (1989): 351 (ecology, as “Amietina n. sp.”);

Davis et al. (2008): 216 (systematics, natural history);

#### Type locality

Uganda, W. Pr., Kibale Forest [Kibale Forest, Western Region, Uganda].

#### Material examined (2♂♂). UGANDA

W Pr, Kibale forest, Ngogo, 6.iv.1984, M. Nummelin *legit* [in MZH].

##### Diagnosis

This species belongs to the *A. larrochei*-group. It can be distinguished from *A. eburnea* and *A. nyassalandica* by male genitalia and by the combination of the lateral margin of genae evenly rounded and the antero-ventral carina of the profemur entire. The latter two characters are also present in *A. larrochei*, which however has slightly finer pronotal punctures. Branco (1989) stated that another character is the double punctuation of the pronotum of *A. larrochei* (*i*.*e*., with larger punctures intermixed with few smaller ones) *vs*. homogeneously punctated in *A. cambeforti*, and the clypeal margin slightly more strongly sinuated next to clypeogenal sutures in *A. larrochei*. It seems to us that these two characters have some degree of variation, so they may not be fully reliable. Anyways, male genitalia and distribution allow to tell the two species apart quite confidently. See also the identification key provided below.

#### Distribution

Uganda.

### *Amietina eburnea* Cambefort, 1981

(Figs. 2B, 4C,L, 5B,E)

Cambefort (1981): 143 (original description);

Cambefort (1984): 106 (ecology);

Cambefort (1985): 339 (faunistics);

Branco (1989): 201 (taxonomy);

Cambefort and Walter (1991): 405 (ecology);

Cambefort and Bordat (2003): 559 (ecology);

Davis et al. (2008): 216 (systematics, natural history);

Moretto (2010a): 469 (faunistics);

#### Type locality

Côte d’Ivoire, Lamto, forêt-galerie du Bandama [gallery forest of Bandama River at Lamto, Ivory Coast].

#### Material examined (3♂♂, 2♀♀). LIBERIA

Mount Nimba, grassfield, 16-25.ix.1979, lowland forest, 500m, fish trap, I. Hanski *legit* [in MZH].

##### Diagnosis

This species belongs to the *A. larrochei*-group. It can be easily separated from similar taxa by the combination of the lateral genal margin evenly rounded and the antero-ventral carina of the profemur interrupted by a row of setigerous punctures, as well as male genitalia. See (Branco, 1989) and the identification key below for further details.

#### Distribution

Ivory Coast, Guinea, Liberia.

### *Amietina larrochei* Cambefort, 1981

(Figs. 2A, 4A–B, J–K, 5A, 6E)

Cambefort (1981): 143 (original description);

Branco (1989): 201 (taxonomy);

Cambefort and Walter (1991): 408 (ecology);

Cambefort and Bordat (2003): 559 (faunistics);

Davis et al. (2008): 216 (systematics, natural history);

Moretto (2010a): 469 (faunistics);

Branco (2011): 11 (nomenclature);

#### Type locality

Cameroun, Mt. Kala [Mount Kala, Centre Region, Cameroon].

#### Material examined (5♂♂, 2♀♀, 1 unsexed specimen). CAMEROON

Mount Kupe, 4.8213 9.7064, 1501m, 29.xii.2015, siftCM12, V. Grebennikov *legit* [1♂, in MZH]; Mount Cameroon, 4.1175 9.0718, 1079m, 28.xii.2015, siftCM07, V. Grebennikov *legit* [1♀, in MZH, with voucher code STL-10208; 1♂, in GMOC]. **LIBERIA**: Mount Nimba, grassfield, 16-25.ix.1979, lowland forest, 500m, fish trap, I. Hanski *legit* [3♂♂, 1♀, 1 unsexed specimen, in MZH].

#### Diagnosis

This species belongs to the *larrochei*-group. See the diagnosis of *A. cambeforti* and the identification key below for further details.

#### Distribution

Cameroon, Gabon, Liberia.

### *Amietina nyassalandica* Branco, 1989

(Figs. 2D, 4D)

Branco (1989): 201 (original description);

Davis et al. (2008): 216 (systematics, natural history);

Josso and Prévost (2008): 2 (taxonomy);

#### Type locality

Malawi, Viphya Mts., Chamambe Forest [Chamambo Forest in the South Viphya Plateau, Malawi].

#### Material examined (1♀). MALAWI

Nyika N.P., 15 km N Thazima Gate, 15.xii.2006, dead millipede, Josso, Juhel & Monfort *legit* [JFJC].

#### Diagnosis

This species belongs to the *larrochei*-group and can be easily distinguished from all other species by the distinctly angular lateral margin of genae and the antero-ventral carina of profemur restricted to the first third of the profemur. See Branco (1989) and the identification key below for further details.

#### Description of the previously unknown female

Similar to the male, except for the following features. **Length:** 4.2 mm. **Head**. Clypeal teeth less prominent, not strongly upturned. **Legs**. Protibia without uncus.

#### Biology

*A. nyassalandica* was found in forest and, in one occasion, attracted to millipedes. It likely has rather generalist feeding habits, similarly to the other species.

#### Distribution

Malawi.

### *Amietina grebennikovi* n. sp

(Figs. 3C, 6B,H)

#### Type locality

-7.845 36.8761, Udzungwa Mountains, Tanzania.

#### Material examined. Holotype (♂)

Tanzania, Udzungwa Mts., −7.845 36.8761, 454m, 6.x.2014, sift2, V. Grebennikov *legit*. **Paratype (♂):** same data as holotype [preserved in 96% ethanol, in MZH, catalogue number KR.41536].

#### Holotype labelling

TANZANIA, Udzungwa Mts. / −7.845 36.8761, 454m / 6.x.2014, sift2 / V. Grebennikov legit [wc, pb] // http://id.luomus.fi / KR.41535 / Univ. of Helsinki / LUOMUS, 2026 [wc, pb, with QR code on the other side] // *Amietina grebennikovi* / Montanaro & Tarasov / n. sp. / HOLOTYPE (♂) / G. Montanaro des. 2026 [rc, pb].

#### Holotype repository

MZH.

##### Diagnosis

*Amietina grebennikovi* belongs to the *mazumbaiensis*-group. See the diagnosis of *A. praedicta* and the identification key below for comparisons with similar species.

#### Description of the holotype (♂)

- male organism, length = 3.2, unit: millimeter;
- male organism, insect region of cuticle
  – insect region of cuticle: ovate;
  – insect region of cuticle: black;
  – insect region of cuticle: glistening;
  – insect region of cuticle encircles anatomical collection has member cuticular seta: light yellow;
- male organism, insect head
  – insect head: pentagonal;
  – insect head, anterior region: attenuate;
  – insect head, clypeus, anterior margin
    * anterior margin, cuticular tooth: bilaterally paired;
    * anterior margin, obtusely notched: present;
  – insect head, antero-lateral margin
    * antero-lateral margin: curved;
    * antero-lateral margin adjacent to genoclypeal sulcus;
  – insect head, anatomical collection has member cuticular puncture;
  – insect head, anterior region
    * anterior region: rugose;
    * anterior region, anatomical collection
      - anatomical collection has member cuticular puncture;
      - anatomical collection: interpunctural distance;
  – insect head, frons, cuticular tubercle
    * cuticular tubercle: arched;
    * cuticular tubercle: obtusely angular;
    * cuticular tubercle coincident with anatomical line
      - anatomical line: transverse orientation;
      - anatomical line coincident with anterior margin of eye cuticle of male organism
  – insect head, antenna has 8 antennomere;
- male organism, pronotum
  – pronotum: trapezoid;
  – pronotum, posterior margin adjacent to cuticular groove of pronotum
  – pronotum, postero-lateral margin: curved;
  – pronotum, anatomical collection
    * anatomical collection has member simple setigerous cuticular puncture: diameter
      - diameter of simple setigerous cuticular puncture
      - diameter larger than diameter of simple setigerous cuticular puncture;
    * anatomical collection, interpunctural distance = 3, unit: diameter of simple setigerous cuticular puncture;
  – pronotum, anatomical collection has member simple setigerous cuticular puncture
- male organism, elytron
  – elytron: width larger than width of pronotum
  – elytron, anatomical collection has member elytral stria
    * elytral stria has element count 8;
    * elytral stria: increased depth;
    * elytral stria, simple setigerous cuticular puncture: diameter larger than width of elytral stria
  – elytron, elytral disc, anatomical collection has member elytral interstria has 2 anatomical row;
- male organism, metathorax, scutellum: absent;
- male organism, hind wing: present;
- male organism, abdominal tergite VIII
  – abdominal tergite VIII: glistening;
  – abdominal tergite VIII, anatomical collection
    * anatomical collection has member simple setigerous cuticular puncture: increased depth;
    * anatomical collection, interpunctural distance = 1, unit: diameter of simple setigerous cuticular puncture;
- male organism, hypomeron
  – hypomeron, anterior region
    * anterior region: concave;
    * anterior region, cuticular carina
      - cuticular carina: curved;
      - cuticular carina: lateral orientation;
  – hypomeron, posterior region, cuticular carina: absent;
- male organism, protibia
  – protibia, distal margin
    * distal margin: oblique orientation;
    * distal margin, uncus: present;
  – protibia has 3 cuticular tooth;
- male organism, profemur, antero-ventral region, cuticular carina: continuous;
- male organism, parameres
  – parameres: elongated;
  – parameres: symmetrical;
  – parameres, postero-ventral region
    * postero-ventral region: sharp;
    * postero-ventral region: ventral orientation;
  – parameres, ventral margin: concave has modifier decreased magnitude;
  – parameres, length = 0.94, unit: length of phallobase;
- male organism, endophallus
  – endophallus, lamella copulatrix has 3 endophallite;
  – endophallus has 3 raspula;

#### Female

Unknown

#### Variation

No significant variation was observed.

#### Etymology

We are glad to dedicate this new species to its collector, the colleague Vasily V. Grebennikov, who has been sampling arthropods extensively in the Eastern Arc Mountains using soil sifting. His efforts with this challenging technique have already disclosed extremely interesting dung beetles, and inspired GM to look for more.

#### Biology

The new species was collected by sifting litter in the wet forest at the foothills of the Udzungwa Mountains, of which it may be endemic. Similarly to other *Amietina*, it may have non-strictly coprophagous habits such as saprophagy and/or necrophagy.

#### Distribution

Tanzania (Udzungwa Mountains).

### *Amietina mazumbaiensis* n. sp

(Figs. 3D–E, 4G–H,M, 6C,G)

#### Type locality

Mazumbai Forest, West Usambara Mountains, Tanzania.

#### Material examined. Holotype (♂)

Tanzania, West Usambara Mts., Mazumbai forest reserve, pitfall, 1500m, 26.xi–2.xii.1996, O. Biström, M. Nieminen, J. Terhivuo & P. Vilkamaa *legit*.

#### Paratype (♀)

same data as the holotype [in MZH, catalogue number GAC.37186].

#### Holotype labelling

TANZANIA W-Usambara / Mts., Mazumbai for. res. / pitfall, 1500 m / 26.XI-2.XII.1996 [wc, pb] // Exp. O.Biströ m, / M.Nieminen, J.Terhivuo / & P.Vilkamaa 1996 // http://id.Luomus.fi / GAC.37187 / Univ. of Helsinki / LUOMUS, 2023 [wc, pb, with QR code on the other side] // *Amietina mazumbaiensis* / Montanaro & Tarasov / n. sp. / **HOLOTYPE (♂)** / G.

Montanaro des. 2026 [rc, pb].

#### Holotype repository

MZH

#### Diagnosis

*Amietina mazumbaiensis* belongs to the *mazumbaiensis*-group together with the closely allied *A. grebennikovi* and *A. praedicta*. Males can be separated from *A. grebennikovi* and *A. praedicta* by: 1) the shape of parameres, which have a higher sclerotised surface below the membranous dorsodistal part in *A. praedicta*; 2) the shape of the head tubercle, which is transverse and straight in *A. praedicta* (Fig. 4E), transverse and slightly ogival in *A. grebennikovi* (Fig. 3C), and less wide and feebly bilobed in *A. mazumbaiensis* (Fig. 4G); 3) the shape of the ventral parameral margin, which is very broadly concave in *A. praedicta* and *A. grebennikovi* (Figs. 6A–B), while it is strongly concave, almost semicircular in *A. mazumbaiensis* (Fig. 6C); 4) the number of raspulae (2 in *A. praedicta* and *A. mazumbaiensis* (Figs. 6A,C), 3 in *A. grebennikovi* (Fig. 6B)); 5) the shape of the lamella copulatrix (Figs. 6G–I). Males of *A. bondarevi* can be readily distinguished from the species of the *mazumbaiensis*-group by the presence of a short head carina (Fig. 4I) and the different male genitalia, with parameres much shorter (0.66 times the length of phallobase instead of 0.92–0.94) and a differently shaped lamella copulatrix (Fig. 6D).

#### Description of the holotype (♂)

- male organism, length = 3.7, unit: millimeter;
- male organism, insect region of cuticle
  – insect region of cuticle encircles anatomical collection has member cuticular seta: light yellow;
  – insect region of cuticle: ovate;
  – insect region of cuticle: black;
  – insect region of cuticle: glistening;
- male organism, insect head
  – insect head, anterior region: attenuate;
  – insect head, clypeus, anterior margin
    * anterior margin, cuticular tooth: bilaterally paired;
    * anterior margin, obtusely notched: present;
  – insect head, antero-lateral margin
    * antero-lateral margin adjacent to genoclypeal sulcus;
    * antero-lateral margin: curved;
  – insect head, anatomical collection has member cuticular puncture;
  – insect head, anterior region
    * anterior region, anatomical collection
      - anatomical collection has member cuticular puncture;
      - anatomical collection: interpunctural distance;
    * anterior region: rugose;
  – insect head, frons, cuticular tubercle
    * cuticular tubercle coincident with anatomical line
      - anatomical line coincident with anterior margin of eye cuticle of male organism
      - anatomical line: transverse orientation;
    * cuticular tubercle: bilobed;
  – insect head, antenna has 8 antennomere;
  – insect head: pentagonal;
- male organism, pronotum
  – pronotum, posterior margin adjacent to cuticular groove of pronotum
  – pronotum, postero-lateral margin: curved;
  – pronotum, anatomical collection
    * anatomical collection, interpunctural distance = 3, unit: diameter of simple setigerous cuticular puncture;
    * anatomical collection has member simple setigerous cuticular puncture: diameter
      - diameter larger than diameter of simple setigerous cuticular puncture;
      - diameter of simple setigerous cuticular puncture
  – pronotum, anatomical collection has member simple setigerous cuticular puncture
  – pronotum: trapezoid;
- male organism, elytron
  – elytron, anatomical collection has member elytral stria
    * elytral stria has element count 8;
    * elytral stria, simple setigerous cuticular puncture: diameter larger than width of elytral stria
    * elytral stria: increased depth;
  – elytron, elytral disc, anatomical collection has member elytral interstria has 2 anatomical row;
  – elytron: width larger than width of pronotum
- male organism, metathorax, scutellum: absent;
- male organism, hind wing: present;
- male organism, abdominal tergite VIII
  – abdominal tergite VIII, anatomical collection
    * anatomical collection, interpunctural distance = 1, unit: diameter of simple setigerous cuticular puncture;
    * anatomical collection has member simple setigerous cuticular puncture: increased depth;
  abdominal tergite VIII: glistening;
- male organism, hypomeron
  – hypomeron, anterior region
    * anterior region, cuticular carina
      - cuticular carina: curved;
      - cuticular carina: lateral orientation;
    * anterior region: concave;
  – hypomeron, posterior region, cuticular carina: absent;
- male organism, protibia
  – protibia has 3 cuticular tooth;
  – protibia, distal margin
    * distal margin, uncus: present;
    * distal margin: oblique orientation;
- male organism, profemur, antero-ventral region, cuticular carina: continuous;
- male organism, parameres
  – parameres, length = 0.93, unit: length of phallobase;
  – parameres, postero-ventral region
    * postero-ventral region: sharp;
    * postero-ventral region: ventral orientation;
  – parameres, ventral margin
    * ventral margin: concave;
    * ventral margin: semicircular;
  – parameres: elongated;
  – parameres: symmetrical;
- male organism, endophallus
  – endophallus has 2 raspula;
  – endophallus, lamella copulatrix has 3 endophallite;

#### Female

Differs from the male by the following features. **Length:** 4.0 mm. **Head**. Head margin more strongly sinuated next to clypeogenal sutures; punctuation of clypeus coarser, denser; tubercle between the eyes absent; frons with a short, arched transverse carina-like bulging (possibly effaced due to wear). **Legs**. Protibia without uncus. **Pygidium**. Less elongated.

#### Variation

Body length is 3.7 mm in the only examined male and 4.0 mm in the only female. The female is rather worn out, as testified by the dull margins of clypeus and protibiae and the very few setae remained on the dorsal body surface. No other significant variation was observed.

#### Etymology

The name refers to the Mazumbai Forest in the West Usambara Mountains, where the new species was collected.

#### Biology

*A. mazumbaiensis* was found in rainforest, where it was collected with pitfall traps. Similarly to other *Amietina*, it may have non-strictly coprophagous habits. It may be endemic to the (West) Usambara Mountains.

### *Amietina praedicta* n. sp

(Figs. 3A–B, 4E–F,N, 6A,I)

#### Type locality

-6.8569 37.6997, Lupanga Peak, Uluguru Mountains, Tanzania

#### Material examined. Holotype (♂)

TZ26-LP3 TANZANIA, Uluguru Mts., Lupanga Peak, −6.8569 37.6997, 1260m, rainforest, 19–21.i.2026, human dung, G. Montanaro *legit* [preserved in 96% ethanol]. **Paratype (♀):** TZ24-A12 TANZANIA, Uluguru Mountains, trail to Bondwa Peak, between −6.8948 37.6711 & −6.8999 37.6720, ca. 1530–1850m, rainforest, millipede trap, 6–9.i.2024, G. Montanaro *legit* [preserved in 96% ethanol, in MZH, catalogue number KR.41538].

#### Holotype labelling

TZ26-LP3 TANZANIA, Uluguru / Mts., Lupanga Peak, −6.8569 / 37.6997, 1260m, rainforest, / 19–21.i.2026, human dung, / G. Montanaro *legit* [wc, pb] // http://id.luomus.fi / KR.41537 / Univ. of Helsinki / LUOMUS, 2026 [wc, pb, with QR code on the other side] // *Amietina praedicta* / Montanaro & Tarasov / n. sp. / HOLOTYPE (♂) / G. Montanaro des. 2026 [rc, pb].

#### Holotype repository

MZH

#### Diagnosis

*A. praedicta* belongs to the *mazumbaiensis*-group. See the diagnosis of *A. mazumbaiensis*

and the identification key provided below for further details.

#### Description of the holotype (♂)

- male organism, insect region of cuticle
  – insect region of cuticle encircles anatomical collection has member cuticular seta: light yellow;
  – insect region of cuticle: ovate;
  – insect region of cuticle: black;
  – insect region of cuticle: glistening;
- male organism, insect head
  – insect head, anterior region: attenuate;
  – insect head, clypeus, anterior margin
    * anterior margin, cuticular tooth: bilaterally paired;
    * anterior margin, obtusely notched: present;
  – insect head, antero-lateral margin
    * antero-lateral margin adjacent to genoclypeal sulcus;
    * antero-lateral margin: curved;
  – insect head, anatomical collection
    * anatomical collection, interpunctural distance = 1.5, unit: diameter of cuticular puncture;
    * anatomical collection has member cuticular puncture;
  – insect head, frons, cuticular tubercle
    * cuticular tubercle coincident with anatomical line
      - anatomical line coincident with anterior margin of eye cuticle of male organism
      - anatomical line: transverse orientation;
    * cuticular tubercle: straight;
    * cuticular tubercle: transverse orientation;
  – insect head, antenna has 8 antennomere;
  – insect head: pentagonal;
- male organism, pronotum
  – pronotum, posterior margin adjacent to cuticular groove of pronotum
  – pronotum, postero-lateral margin: curved;
  – pronotum, anatomical collection
    * anatomical collection, interpunctural distance = 3, unit: diameter of simple setigerous cuticular puncture;
    * anatomical collection has member simple setigerous cuticular puncture: diameter
      - diameter of simple setigerous cuticular puncture
      - diameter larger than diameter of simple setigerous cuticular puncture;
  – pronotum, anatomical collection has member simple setigerous cuticular puncture
  – pronotum: trapezoid;
- male organism, elytron
  – elytron, anatomical collection has member elytral stria
    * elytral stria, simple setigerous cuticular puncture: diameter larger than width of elytral stria
    * elytral stria: increased depth;
    * elytral stria has element count 8;
  – elytron, elytral disc, anatomical collection has member elytral interstria has 2 anatomical row;
  – elytron: width larger than width of pronotum
- male organism, metathorax, scutellum: absent;
- male organism, hind wing: present;
- male organism, abdominal tergite VIII
  – abdominal tergite VIII, anatomical collection
    * anatomical collection, interpunctural distance = 1, unit: diameter of simple setigerous cuticular puncture;
    * anatomical collection has member simple setigerous cuticular puncture: increased depth;
  – abdominal tergite VIII: glistening;
- male organism, hypomeron
  – hypomeron, anterior region
    * anterior region, cuticular carina
      - cuticular carina: curved;
      - cuticular carina: lateral orientation;
    * anterior region: concave;
  – hypomeron, posterior region, cuticular carina: absent;
- male organism, protibia
  – protibia, distal margin
    * distal margin, uncus: present;
    * distal margin: oblique orientation;
  – protibia has 3 cuticular tooth;
- male organism, profemur, antero-ventral region, cuticular carina: continuous;
- male organism, parameres
  – parameres, postero-ventral region
    * postero-ventral region: sharp;
    * postero-ventral region: ventral orientation;
  – parameres, ventral margin: concave has modifier decreased magnitude;
  – parameres: elongated;
  – parameres: symmetrical;
  – parameres, length = 0.92, unit: length of phallobase;
- male organism, endophallus
  – endophallus, lamella copulatrix has 3 endophallite;
  – endophallus has 2 raspula;
- male organism, length = 3.7, unit: millimeter;

#### Female

Differs from the male by the following features. **Body**. Pronotum and head with light greenish bronze hue, with feeble cupreous reflections on the head. **Length:** 3.6 mm. **Head**. Clypeal margin between clypeal teeth less deeply notched; head margin more strongly sinuated next to clypeogenal sutures; punctuation of clypeus coarser, denser; tubercle between the eyes absent; frons with a short, arched transverse carina. **Legs**. Protibia without uncus; protibial teeth slightly wider. **Pygidium**. Less elongated.

#### Variation

Body size is 3.7 mm in the only male examined and 3.6 mm in the only female. The female has a light greenish bronze hue on pronotum and head which is missing from the male, which has just a feeble cupreous hue on the head.

#### Etymology

The specific epithet *praedicta* is a Latin adjective in the feminine form meaning “predicted”. After the initial discovery of *A. mazumbaiensis* in the Usambara Mountains and *A. grebennikovi* in the Udzungwas, GM hypothesized that closely related species may exist in the Uluguru Mountains and other massifs of the Eastern Arc. What a rewarding moment it was when he found *A. praedicta* in the Ulugurus!

#### Biology

*A. praedicta* was found in Afromontane rainforest next to the Bondwa and Lupanga peaks, in the northern part of the Uluguru Mountains Nature Forest Reserve, between 1260 and 1850m a.s.l. It was collected in traps baited with injured millipedes or human dung. It is probably endemic to the Uluguru Mountains.

#### Remarks

The two specimens used for the description were collected in different localities in the northern part of the Ulugurus. We therefore cannot rule out the possibility that the female paratype collected at Bondwa belongs to a different species; however, the substantially identical external morphology (except for sexually dimorphic traits and minor colour differences), the close proximity of the two localities and their similar habitats make us quite confident that the two specimens are conspecific.

#### Distribution

Tanzania (Uluguru Mountains).

### *Amietina bondarevi* n. sp

(Figs. 3F, 4I,O, 6D)

#### Type locality

-6.8306 37.7008, Kigurunyembe (Morogoro), Uluguru Mountains, Tanzania

#### Material examined. Holotype (♂)

TZ24-H2 TANZANIA, Uluguru Mountains, Morogoro, Kigurunyembe, −6.8306 37.7008, ca. 670m, 13–14.i.2024, wet forest, millipede trap, G. Montanaro *legit*.

#### Holotype labelling

TZ24-H2 TANZANIA, Uluguru Mts., / Morogoro, Kigurunyembe, / −6.8306 37.7008, ca. 670m, / 13–14.i.2024, wet forest, / millipede trap, G. Montanaro [wc, pb] // http://id.luomus.fi / KR.41539 / Univ. of Helsinki / LUOMUS, 2026 [wc, pb, with QR code on the other side] // *Amietina bondarevi* / Montanaro & Tarasov / n. sp. / HOLOTYPE (♂) / G. Montanaro des. 2026 [rc, pb].

#### Holotype repository

MZH

#### Diagnosis

*A. bondarevi* is the only species placed in the *bondarevi*-group, characterised by obliquely truncated protibiae (Fig. 4O), the presence of a short carina between the anterior margin of the eyes, as long as 1/3 of the interocular width (Fig. 4I), the irregular pronotal punctuation (Fig. 3F), and the absence of endophallic raspulae (Fig. 6D). The combination of these characters allow an easy distinction from all other described species. Moreover, *A. bondarevi* has a distinct light pinkish cupreous lustre absent from the other species, which have at most a light bronze shine. The parameres and the lamella copulatrix are also highly diagnostic (Fig. 6D). Following the identification key by d’Orbigny (1913), it falls into group 19, section 30(1), but does not match any other described species.

#### Description of the holotype (♂)

- male organism, length = 3.0, unit: millimeter;
- male organism, insect region of cuticle
  – insect region of cuticle encircles anatomical collection has member cuticular seta: light yellow;
  – insect region of cuticle: ovate;
  – insect region of cuticle: pink;
  – insect region of cuticle: glistening;
- male organism, insect head
  – insect head: pentagonal;
  – insect head, anterior region: attenuate;
  – insect head, clypeus, anterior margin
    * anterior margin, cuticular tooth: bilaterally paired;
    * anterior margin, obtusely notched: present;
  – insect head, antero-lateral margin
    * antero-lateral margin adjacent to genoclypeal sulcus;
    * antero-lateral margin: curved has modifier decreased magnitude;
  – insect head, anatomical collection has member cuticular puncture;
  – insect head, antero-medial region, anatomical collection has member cuticular puncture: decreased size;
  – insect head, posterior region: interpunctural distance;
  – insect head, frons, cuticular carina
    * cuticular carina coincident with anatomical line
      - anatomical line connects anterior margin of eye cuticle of left side of insect head
      - anatomical line connects anterior margin of eye cuticle of right side of insect head
    * cuticular carina: straight;
    * cuticular carina: transverse orientation;
  – insect head, antenna has 8 antennomere;
- male organism, pronotum
  – pronotum: trapezoid;
  – pronotum, posterior margin adjacent to cuticular groove of pronotum
  – pronotum, postero-lateral margin: curved;
  – pronotum, anatomical collection
    * anatomical collection, interpunctural distance = 3, unit: diameter of simple setigerous cuticular puncture;
    * anatomical collection has member simple setigerous cuticular puncture: diameter
      - diameter larger than diameter of simple setigerous cuticular puncture;
      - diameter of simple setigerous cuticular puncture
  – pronotum, anatomical collection has member simple setigerous cuticular puncture
- male organism, elytron
  – elytron: width larger than width of pronotum
  – elytron, anatomical collection has member elytral stria
    * elytral stria: increased depth;
    * elytral stria, simple setigerous cuticular puncture: diameter larger than width of elytral stria
    * elytral stria has element count 8;
  – elytron, elytral disc, anatomical collection has member elytral interstria, anatomical row has member simple setigerous cuticular puncture;
- male organism, metathorax, scutellum: absent;
- male organism, hind wing: present;
- male organism, abdominal tergite VIII
  – abdominal tergite VIII: glistening;
  – abdominal tergite VIII, anatomical collection
    * anatomical collection, interpunctural distance = 1, unit: diameter of simple setigerous cuticular puncture;
    * anatomical collection has member simple setigerous cuticular puncture: increased depth;
- male organism, hypomeron
  – hypomeron, anterior region
    * anterior region: concave;
    * anterior region, cuticular carina
      - cuticular carina: curved;
      - cuticular carina: lateral orientation;
  – hypomeron, posterior region, cuticular carina: absent;
- male organism, protibia
  – protibia has 3 cuticular tooth;
  – protibia, distal margin
    * distal margin: oblique orientation;
    * distal margin, uncus: present;
- male organism, profemur, antero-ventral region, cuticular carina: continuous;
- male organism, parameres
  – parameres, length = 0.66, unit: length of phallobase;
  – parameres: shortened;
  – parameres: symmetrical;
  – parameres, postero-ventral region
    * postero-ventral region: sharp;
    * postero-ventral region: ventral orientation;
  – parameres, ventral margin: concave has modifier decreased magnitude;
  – parameres, postero-ventral region: protruding;
- male organism, endophallus
  – endophallus, raspula: absent;
  – endophallus, lamella copulatrix has 2 endophallite;

#### Female

Unknown.

#### Etymology

Named after Vadim Bondarev, in recognition of his contributions to sorting beetle specimens and his assistance with entomological research.

#### Biology

*A. bondarevi* was found in a low elevation disturbed rainforest on the foothills of the northern Uluguru Mountains. It was collected in a trap baited with injured millipedes.

#### Distribution

Tanzania (Uluguru Mountains).

### Identification key to *Amietina* species (translated and modified from Branco (1989))

1. Pronotal punctuation composed of larger punctures intermixed with many distinctly smaller points (Fig. 3). Male and female distal protibial margin oblique relative to main tibial axis (Figs. 4M–O). Male head with a tubercle or a short carina between the anterior margins of the eyes (Figs. 4E,G,I). Female with a short, curved frontal carina (Figs. 4F,H) (females not known in *A. grebennikovi* and *A. bondarevi*) **2**
  – Pronotal punctuation composed of punctures having uniform to irregular sizes, sometimes with micropunctures or a few scattered smaller punctures, but never abundantly covered in both small and distinctly larger punctures (Fig. 2). Male and female distal protibial margin perpendicular to main tibial axis (Fig. 4L), except in *A. larrochei*, in which it is oblique or almost so (Fig. 4J). Male and female head without frontal carina, at most with a transverse bulge (Figs. 4A–D) **4**
2. Male head with a transverse carina placed between the anterior margins of the eyes and occupying approximately 1/3 of interocular distance (Fig. 4I). Integument with light pinkish cupreous lustre. Endophallus without raspulae; male genitalia as in Fig. 6C. Distribution: Tanzania (Uluguru Mountains) ***bondarevi* n. sp**.
  – Male head with a transverse tubercle placed between the anterior margins of the eyes and occupying much less than 1/3 of interocular distance (Figs. 4E,G). Integument without cupreous lustre, occasionally with light bronze lustre. Endophallus with raspulae; male genitalia different **3**
3. Male head tubercle small, slightly bilobed (Fig. 4G). Male head punctuation very coarse (Fig. 4G). Female head carina almost completely effaced (may be due to wear or the only known specimen) (Fig. 4H). Parameral ventral margin strongly concave, almost semicircular in lateral view, apex turned laterally in dorsal view; endophallus with 2 distinct raspulae (Fig. 6C). Lamella copulatrix as in Fig. 6G. Distribution: Tanzania (West Usambara Mountains) ***mazumbaiensis* n. sp**.
  – Male head tubercle slightly transverse, ogival. Male head punctuation very coarse (similar to Fig. 4G). Parameral ventral margin feebly concave in lateral view, apex not turned laterally in dorsal view; endophallus with 3 distinct raspulae (Fig. 6B). Lamella copulatrix as in Fig. 6H. Distribution: Tanzania (Udzungwa Mountains) ***grebennikovi* n. sp**.
  – Male head tubercle slightly transverse, straight (Fig. 4E). Male head punctuation less coarse (Fig. 4E). Female head carina conspicuous (Fig. 4F). Parameral ventral margin feebly concave in lateral view, apex turned laterally in dorsal view; endophallus with 2 distinct raspulae (Fig. 6A). Lamella copulatrix as in Fig. 6I. Distribution: Tanzania (Uluguru Mountains) ***praedicta* n. sp**.
4. Lateral margin of genae angled (Fig. 4D). Anteroventral profemoral carina restricted to the proximal 1/3 of the profemur. Pronotal integument with a few micropunctures intermixed with larger punctures. Male genitalia as in Fig. 5D. Distribution: Malawi ***nyassalandica*** Branco, 1989
  – Lateral margin of genae evenly rounded (Fig. 4A–C). Anterior profemoral carina complete or interrupted by large punctures. Pronotal integument without micropunctures, but sometimes with punctures of different size **5**
5. Anterior profemoral carina interrupted in the middle by a row of large setigerous punctures. Male genitalia as in Figs. 5B,E and 6F. Distribution: Ivory Coast, Guinea, Liberia. ***eburnea*** Cambefort, 1981
  – Anterior profemoral carina complete **6**
6. Distal margin of male and female protibia slightly to clearly oblique with respect to the protibial axis (Figs. 4J–K). Pronotal disc with a few small punctures intermixed with bigger ones. Male genitalia as in Figs. 5A and 6E. Distribution: Cameroon, Gabon, Liberia***larrochei*** Cambefort, 1981
  – Distal margin of male and female protibia perpendicular to the protibial axis (similar to Fig. 4L). Pronotal disc usually without small punctures (although there is some variation). Male genitalia as in Fig. 4C. Distribution: Uganda ***cambeforti*** Branco, 1989

## DISCUSSION

### Semantic publishing workflow

The need to publish taxonomic papers as structured text has long been recognized (Penev et al., 2010, 2011; Miller et al., 2012). In conventional documents, taxon names, treatments, specimen records, and figures are visible to readers but are not always identified in a way that computers can easily interpret. Structured markup gives these elements explicit roles, allowing them to be found, linked, extracted, and reused. TaxPub format introduced such markup for taxonomic literature, while Pensoft’s ARPHA platform allows manuscripts to be written and reviewed online and published as structured XML and HTML (Penev et al., 2011; Stoev, 2018). Together with persistent identifiers, accessible repositories, and suitable metadata, these developments can make taxonomic publications more consistent with the FAIR principles (Wilkinson et al., 2016).

Comparable systems have been developed outside taxonomy. Pandoc Scholar generates several publication formats from one plain-text source, whereas Manubot combines text-based writing, Git version control, and automated manuscript generation (Krewinkel and Winkler, 2017; Himmelstein et al., 2019). Both follow the same principle: content should be recorded once in a structured source and reused to produce the required outputs.

Our workflow applies this principle to semantic species descriptions, an emerging approach in taxonomy (Balhoff et al., 2013; Deans et al., 2015; Montanaro et al., 2024a). Phenoscript generates computable phenotype data and readable text from the same source. GitHub records changes, and Overleaf imports the generated Markdown into the LATEX manuscript. A correction is therefore made in Phenoscript and propagated to the manuscript after regeneration and synchronization, rather than copied into several versions by hand. This keeps the published treatment aligned with its semantic source, and preserves a shared revision history (Ram, 2013; Braga et al., 2023).

Wider adoption of these principles in taxonomy faces two main barriers. First, LATEX remains uncommon among taxonomists and is not accepted by many taxonomic journals. Yet its use can reduce manual reformatting and streamline production, potentially lowering publication costs (Bar and Wang, 2021; Krewinkel and Winkler, 2017). Broader support for LATEX among authors, journals, and publishers would therefore benefit taxonomy.

Second, Phenoscript requires authors to learn its syntax and the basic principles of ontologies. Wider use will depend on community-led training and engagement. Artificial-intelligence tools may help convert existing descriptions or prepare initial Phenoscript drafts, although their output will still require validation and expert review.

A smaller practical limitation is that some Overleaf features require a subscription, although institutional access may be available. Overleaf is convenient but not essential: authors can instead use a free local LATEX distribution and text editor together with GitHub. The essential requirement is a text-based, version-controlled manuscript into which generated Phenoscript outputs can be incorporated without duplication.

This study shows that semantic descriptions can be used throughout a complete taxonomic revision. Publication alone, however, does not make them readily available for comparative use. A centralized repository is badly needed where semantic phenotypes can be deposited, retrieved, queried, and compared across taxa and publications. Besides providing these services, such a repository could also provide a shared environment for engaging the taxonomic community in the adoption and development of new technologies.

### *Amietina* microendemism in the Eastern Arc Mountains

The Eastern Arc Mountains in Tanzania and Kenya are one of the most important hotspots of biodiversity on the Earth, hosting an exaggerated amount of endemic plant and animal species (Burgess et al., 2007). However, dung beetle diversity in the EAMs is virtually unknown. The only available study on the subject reported an instance of microallopatric speciation within a single mountain block in the epactoidine genus *Grebennikovius* Mlambo, Sole & Scholtz, 2019 (Montanaro et al., 2024b), but nothing is known about endemism of other dung beetle lineages. The *Amietina* species described here are likely endemic to the separate mountains where they were discovered. In particular, *A. grebennikovi, A. mazumbaiensis*, and *A. praedicta* form a complex of morphologically very similar species that may have originated through allopatric speciation in the Udzungwa, West Usambara, and Uluguru Mountains, respectively, making them the first documented case of microendemism in which separate EAM forests host different, strictly related dung beetle species. This suggests that additional *Amietina* species occur in other EAM regions, including the Rubeho Mountains, where a putative *Amietina* specimen was collected (see Methods) but could not be examined in the present study. An ongoing survey shows that this pattern is probably widespread across EAMs dung beetles, whose fauna is largely undescribed (unpublished data) and will be the target of future contributions.

## CONCLUSIONS

Taxonomy must keep pace with the rapid development of digital biology. Although analytical methods have advanced considerably, species descriptions and manuscripts are still often prepared as conventional text, limiting efficient workflows, computational reuse, and integration with other biodiversity data.

This study demonstrates that semantic species descriptions can be incorporated into a complete taxonomic revision and carried directly from their computable source into a publication-ready manuscript. The integrated use of Phenoscript, GitHub, and Overleaf provides a practical model for producing taxonomic treatments that remain readable to taxonomists while being available for computational reuse.

The revision also reveals previously undocumented diversity in the Eastern Arc Mountains by adding four new species of *Amietina*. Their restricted distributions provide further evidence that separate mountain blocks harbor distinct dung beetle faunas and emphasize the importance of documenting the remaining rainforest fragments. Together, these methodological and taxonomic results show how structured digital approaches can support both the study of biodiversity and the preservation of the knowledge derived from it.

## ACKNOWLEDGEMENTS

We are indebted to: Vasily V. Grebennikov (Canadian Food Inspection Agency, Ottawa, ON, Canada) for entrusting us with the invaluable dung beetle material collected on the EAMs; Jean-François Josso (Muzillac, France) for allowing us to study the female of *Amietina nyassalandica*; Jaakko Mattila, Lotta Leinonen, Olof Biström and Pekka Vilkamaa (MZH) for their help in Helsinki; Philippe Moretto (Toulon, France) and Fernando Vaz-de-Mello (Universidade Federal de Mato Grosso, Cuiabá, Brasil) for sharing their scarab bibliography. We thank the Tanzania Wildlife Research Institute (TAWIRI) and the Tanzania Commission for Science and Technology (COSTECH) fo releasing the necessary research permits. GM is sincerely grateful to the Tanzanian authorities and collaborators that helped obtaining permits and developing a project on EAMs dung beetles, including Alain Pauly (TAWIRI) and Prof. Abdallah Jumanne and Maulid Mwatawala (Sokoine University of Agriculture). This study was supported by the Research Council of Finland (grants 361983, 346294, and 369299); field work was supported by the Linnean Society of London’s Percy Sladen Memorial Fund 2025, awarded to GM.

